# Immune aging captures complementary aging biology beyond epigenetic clocks

**DOI:** 10.64898/2026.05.19.726183

**Authors:** Kohav Tal-Porath, Timothy J. Few-Cooper, Shai S. Shen-Orr

**Affiliations:** Department of Immunology, Rappaport Faculty of Medicine, Technion – Israel Institute of Technology, Haifa, Israel

## Abstract

Biological aging clocks are typically evaluated through competitive benchmarking, implicitly assuming that a single metric can sufficiently capture the complexities of aging^1-6^. Here, we tested an alternative hypothesis: that distinct clock types capture orthogonal dimensions of aging and therefore yield greater value when integrated. Using the Framingham Heart Study, we compared the immune-aging metric, IMM-AGE, with established DNA methylation clocks and found that integrated models consistently outperformed single-clock approaches. To investigate the basis of this complementarity, we derived IMMAGE-Epi, a 22-CpG methylation surrogate of IMM-AGE which exhibited minimal overlap with canonical epigenetic clock CpGs, suggesting that immune aging is associated with a distinct methylomic feature and pathway space rather than representing a reformulation of existing clock architectures. Together, our findings support an emerging multidimensional model of biological aging in which integrating orthogonal biological clocks may offer greater translational utility than competitive single-clock optimization.

## Introduction

Chronological age remains central to clinical decision-making despite marked heterogeneity in biological aging trajectories among individuals of the same age. Biological clocks have emerged to address this limitation by quantifying aging using molecular and physiological biomarkers, with DNA methylation clocks becoming one of the most prominent translational candidates owing to their reproducibility, accessibility, and strong associations with mortality and age-associated disease^7-13^. However, the field has increasingly adopted a benchmark-driven framework in which each new clock is positioned primarily in terms of statistical superiority over its predecessors^1,2^. Such framing may be conceptually limiting. Aging is a distributed, multi-system process shaped by genetics, environmental exposures, immune history, metabolism and stochastic biological drift^3-5,14^. Different measurement systems may therefore be expected to capture distinct dimensions of aging biology rather than redundant representations of the same latent construct^6,15^.

IMM-AGE was developed as an unsupervised immune-aging trajectory derived from high-dimensional immune profiling, independent of chronological-age supervision^16^. Unlike canonical epigenetic clocks, IMM-AGE captures immune system remodeling directly rather than through methylation correlates of age or mortality. Therefore, here, we tested whether IMM-AGE and epigenetic clocks should be viewed as competitors or complements [Figure 1A]. Specifically, we asked whether integrating these approaches improves mortality prediction, whether immune aging leaves a distinct methylomic signature, and whether this signature reflects coherent biological programs distinct from canonical epigenetic clock architectures.

**Figure 1.**
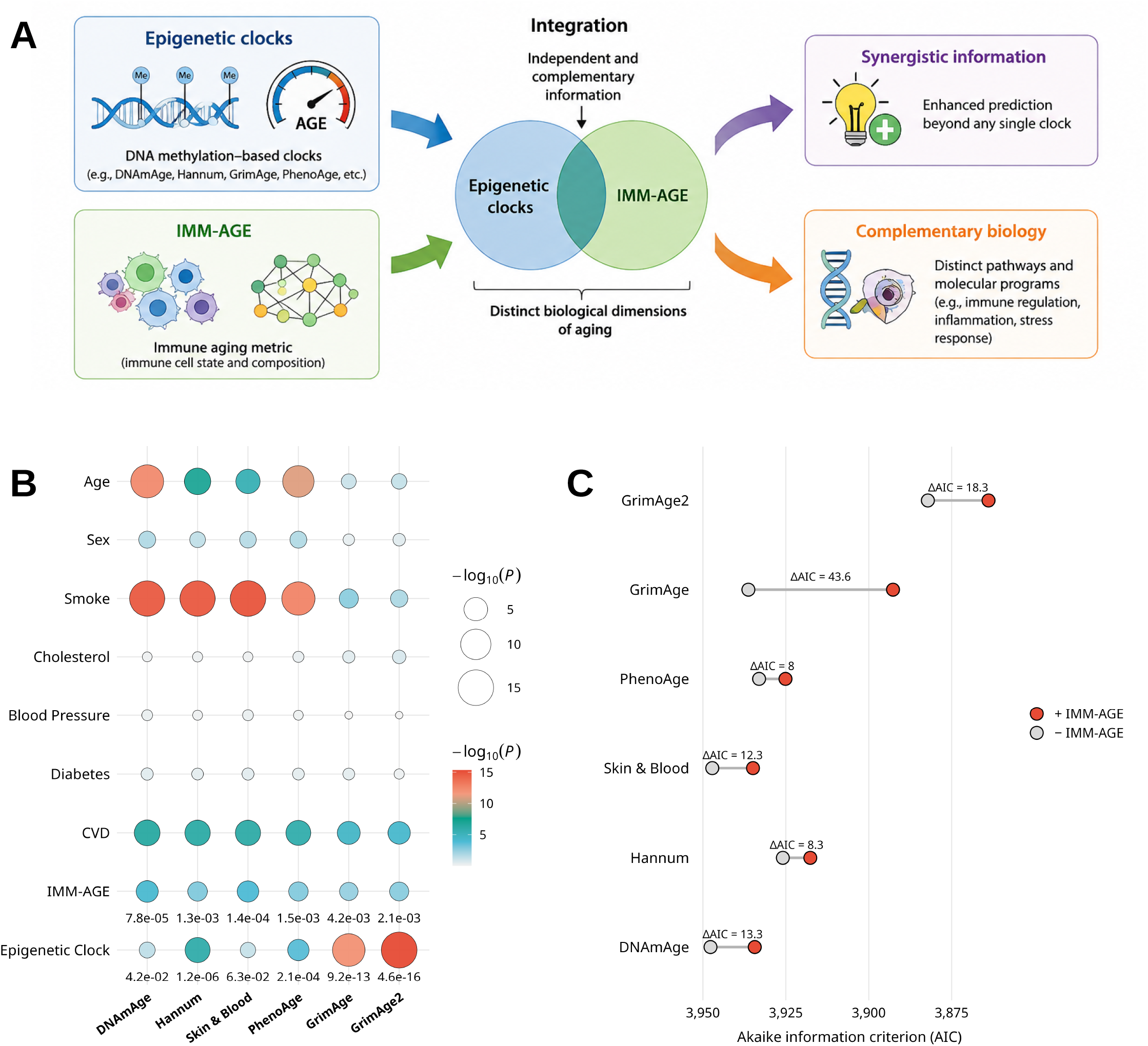
Integration of immune and epigenetic aging clocks improves mortality prediction. (**a**) Conceptual framework illustrating the central hypothesis of the study. Rather than treating biological aging clocks as competing alternatives, we tested whether clocks derived from distinct biological modalities capture complementary dimensions of aging biology. Epigenetic clocks reflect DNA methylation-based aging signals, whereas IMM-AGE captures immune aging state. Integration of orthogonal biological information may therefore improve prediction beyond any single clock alone. (**b**) Bubble significance matrix showing the association of conventional mortality risk covariates, IMM-AGE, and the corresponding epigenetic clock within Cox proportional hazards mortality models containing individual epigenetic clocks. Bubble size and colour indicate statistical significance (™log10(*P*)). IMM-AGE remained independently associated with all-cause mortality across all pairwise models, supporting non-redundant prognostic information relative to canonical epigenetic clocks. (**c**) Akaike information criterion (AIC) comparisons for mortality models with and without IMM-AGE across individual epigenetic clock frameworks. Lower AIC indicates improved model fit. Addition of IMM-AGE consistently improved model performance, with the largest improvement observed when combined with GrimAge2, supporting complementary predictive value rather than substitution.

### Orthogonal biological clocks improve mortality prediction

To test whether IMM-AGE contributes information beyond established epigenetic clocks, we evaluated all-cause mortality associations in the Framingham Heart Study (n=2222) using multivariable Cox regression models containing conventional clinical covariates together with IMM-AGE and individual epigenetic clocks. Remarkably, IMM-AGE remained significantly associated with mortality across all pairwise models, including those containing DNAmAge^7^ (*P* = 7.85 × 10^−5^), Hannum^8^ (*P* = 1.28 × 10^−3^), Skin & Blood^9^ (*P* = 1.4 × 10^−4^), PhenoAge^10^ (*P* = 1.52 × 10^−3^), GrimAge^11^ (*P* = 4.23 × 10^−3^), and GrimAge2^12^ (*P* = 0.0021) [Figure 1B]. As expected, mortality-trained clocks such as GrimAge and GrimAge2 exhibited particularly strong individual predictive performance^11,12^ (*P* =9.21 × 10^−13^, and *P* = 4.61 × 10^−16^ respectively), whereas first-generation chronological-age clocks performed less consistently^7,8^. To further assess incremental utility, we compared nested models with and without IMM-AGE. Consistent with the previous results, addition of IMM-AGE improved model fit across every pairwise comparison and direct model comparison using Akaike information criterion further supported the complementarity of the two clock-types, as the best-performing model consisted of GrimAge2 + IMM-AGE (AIC 3863.9), significantly outperforming GrimAge2 alone (3882.2, ΔAIC = 18.3) [Figure 1C]. Nevertheless, when all biological clocks were forced into direct competition without clinical covariates, IMM-AGE lost nominal significance (*P* = 0.089), while GrimAge2 remained dominant (*P* = 2.03 × 10^−6^, data not shown). However, we do not interpret this as evidence of redundancy. Rather, such a model represents an analytically adversarial setting that maximizes covariance between partially overlapping predictors and is unlikely to reflect practical clinical implementation, where biological markers are integrated with demographic and clinical context. Taken together, these findings suggest that IMM-AGE, as applied in a real-world-like setting, is not simply a weaker competitor within the existing clock hierarchy, but instead captures aging-relevant biological variance not fully represented by epigenetic clocks, therefore supporting a shift from winner-takes-all benchmarking toward integrative modeling of complementary dimensions and axes of aging.

### IMMAGE-Epi reveals distinct immune-aging methylation programs

To investigate the biological basis of this complementarity, we next asked whether immune aging leaves a detectable methylomic signature. Using whole-blood DNA methylation data from the Framingham Heart Study, we derived an epigenetic surrogate of IMM-AGE, termed IMMAGE-Epi. Due to the extremely high dimensionality of methylation data, feature selection was performed in multiple stages. Initial univariate screening identified CpGs significantly associated with IMM-AGE, followed by repeated regularised regression with stability-based feature selection to identify robust predictors. This yielded a sparse 22-CpG methylation signature capable of reconstructing IMM-AGE with strong performance (r = 0.82, P < 2.2 × 10^−16^) [Figure 2A]. These findings indicate that immune aging, as captured by IMM-AGE, is associated with a measurable epigenetic signature. To exclude the possibility that IMMAGE-Epi simply reflects shifts in immune cell composition, we regressed deconvolution-inferred cell-composition variance from the methylation matrix; predictive performance remained substantial (r = 0.73, P < 2.2 × 10^−16^), arguing against simple compositional confounding. We next asked whether IMMAGE-Epi simply reformulates biology already captured by conventional epigenetic clocks. To assess feature-level overlap, we expanded IMMAGE-Epi to include correlated CpGs associated with the model (IMMAGE-Epi-expanded), yielding a feature set of >2,000 loci. Despite using this permissive definition, overlap with established epigenetic clock architectures remained minimal, supporting the molecular, orthgonal distinctiveness of IMMAGE-Epi and, in-turn, IMMAGE [Figure 2B].

**Figure 2.**
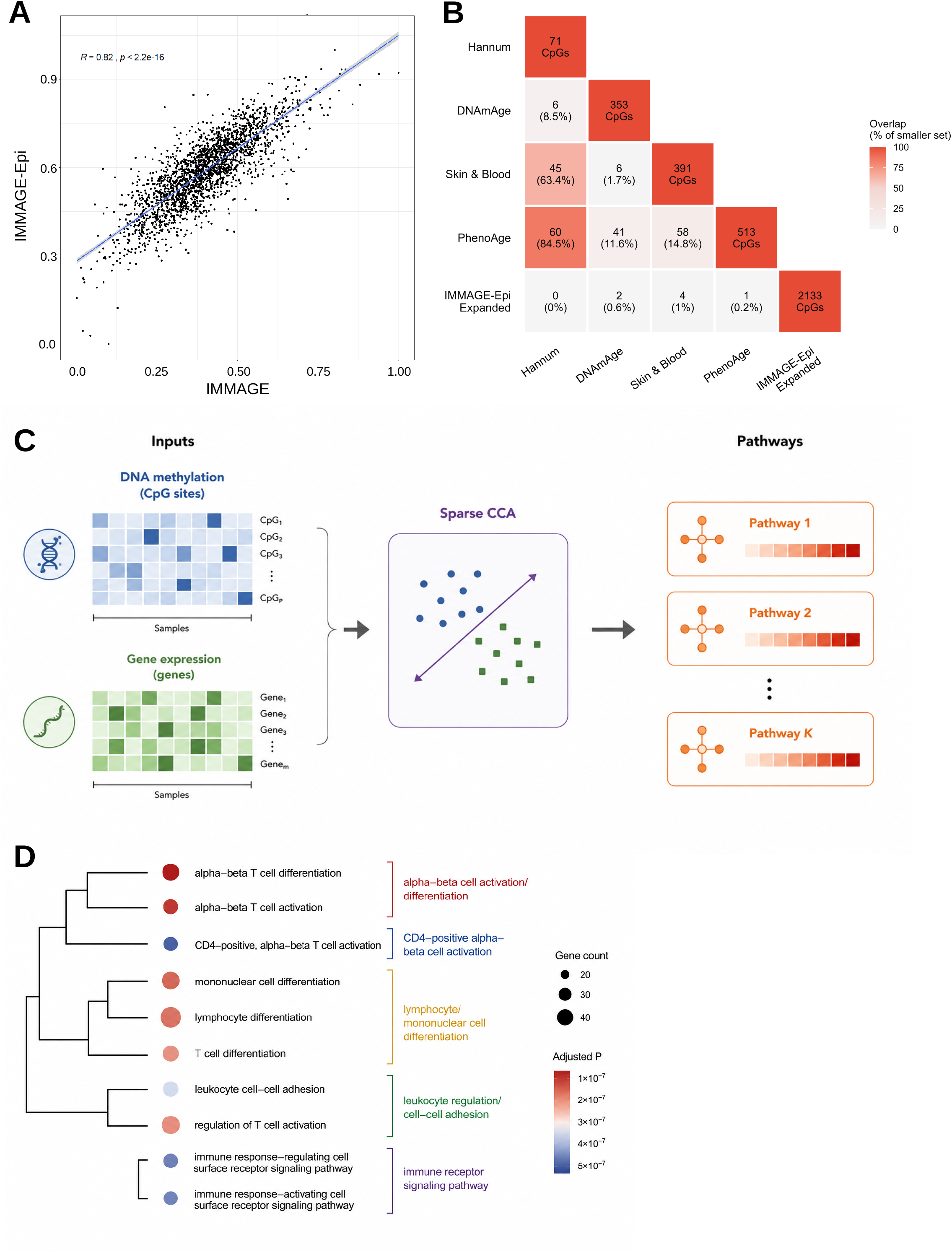
IMMAGE-Epi identifies a distinct methylomic signature of immune aging-related. (**a**) Prediction accuracy of IMM-AGE from DNA methylation data (IMMAGE-Epi). IMM-AGE was strongly predictable from a 22 CpG methylation profile, indicating the presence of a measurable epigenetic correlate of immune aging. Line represents linear regression with 95% confidence interval shading. (**b**) Pairwise overlap between CpG features from IMMAGE-Epi-expanded and established epigenetic clock architectures. To assess molecular overlap more permissively, IMMAGE-Epi was expanded to include correlated CpGs associated with the predictive model. Values indicate absolute overlap counts, with percentages calculated relative to the smaller CpG set in each pairwise comparison. Despite the expanded feature space, overlap with canonical clock architectures remained minimal, supporting the molecular distinctiveness of IMMAGE-Epi. (**c**) Schematic overview of sparse canonical correlation analysis (sparse CCA) used for biological interpretation of IMMAGE-Epi-associated methylation features. DNA methylation and paired gene expression matrices were jointly modelled to identify coupled latent components linking CpG methylation patterns with transcriptional programmes. Gene features contributing to correlated components were subsequently subjected to pathway enrichment analysis. (**d**) Functional enrichment analysis of gene programmes associated with IMMAGE-Epi-linked sparse CCA components. Hierarchical clustering of enriched pathways identified coherent immune regulatory modules, including T cell activation and differentiation, leukocyte regulation, and immune receptor signalling pathways. Dot size indicates the number of contributing genes; dot colour indicates adjusted *P* value. These findings support a biologically coherent interpretation of IMMAGE-Epi as capturing immune regulatory ageing biology rather than simple compositional effects.

While epigenetic clocks often achieve strong predictive performance, biological interpretability remains a major limitation. CpG-based models are frequently interpreted using simplistic proximity-based annotation, in which CpGs are assigned to nearby genes despite often weak or uncertain regulatory relationships^19,20^. To improve interpretability, we linked IMMAGE-Epi-associated methylation programs to transcriptional programs using sparse canonical correlation analysis (sCCA)^21,22^, enabling correlated CpG–gene modules to be identified directly rather than inferred by genomic proximity alone [Figure 2C]. This analysis revealed coherent immune-associated transcriptional modules linked to IMMAGE-Epi methylation features. Rather than reflecting diffuse age-associated methylation drift, the identified programs were enriched for classical immune regulatory processes consistent with immune activation, inflammatory signalling, and immune remodelling [Figure 2D]. These findings support a biologically coherent interpretation of IMMAGE-Epi as an immune-aging-associated regulatory program rather than a purely statistical predictor. Taken together, these findings further support the biological orthogonality of immune-aging and canonical epigenetic clock architectures, suggesting that their predictive complementarity reflects distinct underlying aging biology rather than simple statistical redundancy.

## Discussion

The biological aging field has largely advanced through competitive optimisation, with new clocks typically judged by whether they outperform existing models in predicting age, mortality, or age-related decline^1,2^. Although this framework has accelerated methodological progress, it also reinforces the assumption that biological aging can be adequately captured by a single axis. Our findings support a different view. IMM-AGE was not intended to replace leading epigenetic clocks, nor did it uniformly outperform them. Rather, its value emerged through complementarity: across mortality models, IMM-AGE consistently contributed independent predictive information, with the strongest performance arising from integration rather than substitution. IMMAGE-Epi reinforced the biological basis of this complementarity. The identification of a measurable methylomic signature of immune aging, its persistence after adjustment for immune cell composition, and its minimal overlap with canonical clock architectures together suggest that immune aging occupies a distinct molecular axis of aging rather than recapitulating the processes reflected by existing epigenetic clocks. Sparse canonical correlation analysis further linked these features to coherent immune regulatory programmes, supporting biological orthogonality rather than statistical artefact.

These observations have broader translational implications. Precision medicine is unlikely to converge on a universal biological age score derived from a single molecular modality^6,15,23^. A more plausible framework may involve integrating complementary biomarkers that capture distinct dimensions of aging biology—including immune remodelling, epigenetic change, metabolic state, proteomic variation, and functional resilience^23-25^. While this study has limitations, including analysis within a single cohort and the use of IMMAGE-Epi primarily as an interpretive rather than clinical construct, the central implication is conceptual: progress in biological aging biomarker development may depend less on identifying the single best clock than on determining which clocks, and which combinations of clocks, broadly capture distinct aging-related biology - and how best to integrate them for practical use in the clinic.

## Author Contributions

T.J.F-C and S.S.S-O conceived of the study. K.T.-P led the computational and statistical analyses, generated the figures and wrote the manuscript. T.J.F.-C. contributed data analysis, interpretation of results, and manuscript writing and editing. S.S.S.-O. contributed to interpretation of results, and manuscript writing and editing. T.J.F-C and S.S.S-O jointly supervised the work.

### Author Disclaimers

Shai S. Shen-Orr is a consultant for and holds equity in CytoReason. In addition he is the co-CSO of the Human Immunome Project, where he serves on a voluntary basis.

## Acknowledgments

This research was supported by NIH joint NIAID & NIA award P01AI153559, and the ISRAEL SCIENCE FOUNDATION (grant No. 1626/20), within the Israel Precision Medicine Partnership program.

## Methods

### Study Cohort

Analyses were conducted using data from the Framingham Heart Study (FHS) Offspring Cohort, a longitudinal community-based cohort established to investigate cardiovascular and aging-related phenotypes. Participants with available DNA methylation, transcriptomic and phenotypic data, together with corresponding IMM-AGE estimates, were included (n=2222). Mortality follow-up data were obtained from linked longitudinal outcome records. All analyses were conducted on de-identified data obtained through approved access procedures.

### DNA methylation preprocessing and epigenetic clock calculation

Genome-wide DNA methylation data were generated using the Illumina HumanMethylation450 BeadChip platform. Preprocessing was performed in R (v4.3.1) using the *minfi* package (v1.46.0). Raw methylation intensities underwent background correction, normalisation and quality control according to standard workflows, with methylation quantified as β-values. Established epigenetic clock estimates were calculated using publicly available implementations via the Clock Foundation DNA methylation age calculator, including Horvath DNAmAge, Hannum age, Skin & Blood clock, PhenoAge, GrimAge and GrimAge2. These clocks were used as provided for downstream comparative modelling.

### Mortality modelling

Associations between biological aging measures and all-cause mortality were assessed using multivariable Cox proportional hazards regression implemented in R using the *survival* package. Models included chronological age, sex, smoking status, cholesterol, blood pressure, diabetes and cardiovascular disease as covariates. To evaluate complementarity between IMM-AGE and epigenetic clocks, pairwise models were constructed containing IMM-AGE and a single epigenetic clock alongside conventional covariates. Hazard ratios, Wald statistics and model significance were extracted from fitted models. Model fit was compared using Akaike information criterion (AIC), with lower values indicating improved fit. Nested model comparisons were performed using likelihood ratio testing to assess the incremental contribution of IMM-AGE beyond epigenetic clock models alone.

### Development of IMMAGE-Epi

To derive an epigenetic surrogate of IMM-AGE (IMMAGE-Epi), CpGs associated with IMM-AGE were first identified using univariate linear regression across genome-wide methylation features. Resulting *P* values were adjusted using the Benjamini–Hochberg procedure, and CpGs passing a false discovery threshold of adjusted *P* < 0.05 were retained for model development. A penalised regression model was then trained using the *biglasso* package in R, applying LASSO regularisation with internal cross-validation for feature selection and coefficient optimisation. Data were partitioned into training and held-out testing subsets using a 75:25 split. Model performance was assessed by Pearson correlation between predicted and observed IMM-AGE values in the independent test set. The final IMMAGE-Epi model comprised 22 CpG loci selected through penalised feature reduction.

### Immune cell composition adjustment

To assess whether IMMAGE-Epi primarily reflected variation in immune cell composition rather than immune regulatory biology, immune cell abundances were inferred from paired transcriptomic data using CIBERSORTx with the LM22 leukocyte reference signature matrix. Quantile normalisation and batch correction were enabled, with 100 permutations used for deconvolution.

To account for compositional confounding, principal component analysis was applied to inferred immune cell abundance estimates using *scikit-learn* in Python. Principal components capturing the majority of cell composition variance were regressed from the methylation matrix using *limma* (*removeBatchEffect*), and IMMAGE-Epi modelling was repeated on the adjusted methylation data.

### Definition of IMMAGE-Epi-expanded

To assess broader molecular overlap between IMMAGE-Epi and canonical epigenetic clock architectures, an expanded IMMAGE-Epi feature set was defined by identifying CpGs significantly correlated with the 22-CpG IMMAGE-Epi model features across the methylation dataset. This permissive feature set was used exclusively for overlap analysis and was not used for predictive modelling. Pairwise overlap with established clock CpG architectures was quantified using absolute overlap counts and expressed relative to the smaller CpG set in each comparison.

### Sparse canonical correlation analysis

To investigate the biological programmes associated with IMMAGE-Epi-linked methylation features, sparse canonical correlation analysis (sparse CCA) was performed on paired DNA methylation and transcriptomic data. Sparse CCA identifies correlated latent components between two high-dimensional omics matrices while enforcing sparsity to improve interpretability. Analysis was implemented in Python using the *cca-zoo* framework, using sparse canonical variates to identify coupled methylation–transcriptional programmes. Input methylation features were restricted to IMMAGE-Epi-associated CpGs and their correlated expanded feature set. Transcriptomic inputs consisted of matched gene expression profiles from the same participants. Sparsity penalties were tuned empirically to optimise stable component recovery while maintaining interpretability. Genes with non-zero loadings in significant canonical components were extracted for downstream pathway analysis.

### Pathway enrichment analysis

Functional enrichment analysis was performed in R using *clusterProfiler* with Gene Ontology biological process annotations from *org*.*Hs*.*eg*.*db*. Enrichment testing was conducted on genes identified from sparse CCA-derived canonical components. Multiple testing correction was performed using the Benjamini–Hochberg procedure, and pathways with adjusted *P* < 0.05 were considered significant. Enriched pathways were clustered based on functional similarity for visualisation.

### Statistical analysis and software

Statistical analyses were performed using R (v4.3.1) and Python. R packages included *tidyverse, survival, minfi, biglasso, limma, clusterProfiler, org*.*Hs*.*eg*.*db* and associated dependencies. Python analyses used *pandas, NumPy, scikit-learn, SciPy* and *cca-zoo*. All statistical tests were two-sided unless otherwise stated. Multiple testing correction was performed using the Benjamini–Hochberg procedure where applicable.

## Notes

### Competing Interest Statement

The authors have declared no competing interest.

### Summary of Updates

Include author disclaimers and funding acknowledgements within the main PDF. No other changes.

## References

1. Jylhävä, J., Pedersen, N. L. & Hägg, S. Biological age predictors. EBioMedicine 21, 29–36 (2017).

2. Bell, C. G. et al. DNA methylation aging clocks: challenges and recommendations. Genome Biology 20, 249 (2019).

3. López-Otín, C., Blasco, M. A., Partridge, L., Serrano, M. & Kroemer, G. The hallmarks of aging. Cell 153, 1194–1217 (2013).

4. Kennedy, B. K. et al. Geroscience: linking aging to chronic disease. Cell 159, 709–713 (2014).

5. Ferrucci, L. et al. Measuring biological aging in humans: a quest. Aging Cell 19, e13080 (2020).

6. Justice, J. N. et al. Frameworks for proof-of-concept clinical trials of interventions that target fundamental aging processes. J. Gerontol. A Biol. Sci. Med. Sci. 73, 141–151 (2018).

7. Horvath, S. DNA methylation age of human tissues and cell types. Genome Biology 14, R115 (2013).

8. Hannum, G. et al. Genome-wide methylation profiles reveal quantitative views of human aging rates. Mol. Cell 49, 359–367 (2013).

9. Horvath, S. et al. Epigenetic clock for skin and blood cells applied to Hutchinson Gilford Progeria Syndrome and ex vivo studies. Aging 10, 1758–1775 (2018).

10. Levine, M. E. et al. An epigenetic biomarker of aging for lifespan and healthspan. Aging 10, 573–591 (2018).

11. Lu, A. T. et al. DNA methylation GrimAge strongly predicts lifespan and healthspan. Aging 11, 303–327 (2019).

12. Lu, A. T. et al. DNA methylation GrimAge version 2. Aging 14, 4812–4838 (2022).

13. Hillary, R. F. et al. Multi-method comparison of biological age estimation approaches in the UK Biobank. Nature Communications 12, 5262 (2021).

14. Campisi, J. et al. From discoveries in ageing research to therapeutics for healthy ageing. Nature 571, 183–192 (2019).

15. Belsky, D. W. et al. Quantification of the pace of biological aging in humans through a blood test, the DunedinPACE DNA methylation algorithm. eLife 11, e73420 (2022).

16. Alpert, A. et al. A clinically meaningful metric of immune age derived from high-dimensional longitudinal monitoring. Nature Medicine 25, 487–495 (2019).

17. Newman, A. M. et al. Determining cell type abundance and expression from bulk tissues with digital cytometry. Nature Biotechnology 37, 773–782 (2019).

18. Zhou, W., Laird, P. W. & Shen, H. Comprehensive characterization, annotation and innovative use of Infinium DNA methylation BeadChip probes. Nucleic Acids Research 45, e22 (2017).

19. Roadmap Epigenomics Consortium et al. Integrative analysis of 111 reference human epigenomes. Nature 518, 317–330 (2015).

20. ENCODE Project Consortium. Expanded encyclopaedias of DNA elements in the human and mouse genomes. Nature 583, 699–710 (2020).

21. Witten, D. M., Tibshirani, R. & Hastie, T. A penalized matrix decomposition, with applications to sparse principal components and canonical correlation analysis. Biostatistics 10, 515–534 (2009).

22. Chapman, A. et al. A python package for canonical correlation analysis. J. Open Source Softw. 6, 3823 (2021).

23. Lehallier, B. et al. Undulating changes in human plasma proteome profiles across the lifespan. Nature Medicine 25, 1843–1850 (2019).

24. Ahadi, S. et al. Personal aging markers and ageotypes revealed by deep longitudinal profiling. Nature Medicine 26, 83–90 (2020).

25. Belsky, D. W. et al. Eleven telomere, epigenetic clock, and biomarker-composite quantifications of biological aging: do they measure the same thing? Am. J. Epidemiol. 187, 1220–1230 (2018).

